# PorSignDB: a database of *in vivo* perturbation signatures for dissecting clinical outcome of PCV2 infection

**DOI:** 10.1101/341644

**Authors:** Nicolaas Van Renne, Ruifang Wei, Nathalie Pochet, Hans J. Nauwynck

## Abstract

Porcine Circovirus Type 2 (PCV2) is a pathogen that has the ability to cause often devastating disease manifestations in pig populations with major economic implications. How PCV2 establishes subclinical persistence and why certain individuals progress to lethal lymphoid depletion remain to be elucidated. Here we present PorSignDB, a gene signature database describing *in vivo* porcine tissue physiology that we generated from a large compendium of *in vivo* transcriptional profiles and that we subsequently leveraged for deciphering the distinct physiological states underlying PCV2-affected lymph nodes. This systems biology approach indicated that subclinical PCV2 infections shut down the immune system. A robust signature of PCV2 disease emphasized that immune activation is dysfunctional in subclinical infections, however, in contrast it is promoted in PCV2 patients with clinical manifestations. Functional genomics further uncovered IL-2 as a driver of PCV2-mediated disease and we identified STAT3 as a druggable PCV2 host factor candidate. Our systematic dissection of the mechanistic basis of PCV2 reveals that subclinical and clinical PCV2 display two diametrically opposed immunotranscriptomic recalibrations that represent distinct physiological states *in vivo*, which suggests a paradigm shift in this field. Finally, our PorSignDB signature database is publicly available as a community resource (http://www.vetvirology.ugent.be/PorSignDB/, included in Gene Sets from Community Contributors http://software.broadinstitute.org/gsea/msigdb/contributed_genesets.jsp) and provides systems biologists with a valuable tool for catalyzing studies of human and veterinary disease.

**Author Summary:** Porcine Circovirus Type 2 (PCV2) is a small but economically important pathogen circulating endemically in pig populations. Although PCV2 causes mostly chronic subclinical infections, many individuals develop a lethal form of circoviral disease consisting of a collapse of lymphoid tissue. In order to provide a fresh look at how PCV2 reprograms host tissue, we created PorSignDB, a compendium of hundreds of transcriptomic gene-expression signatures derived from primary porcine tissue specimens of well over 1500 patients or lab animals. By leveraging PorSignDB on transcriptomic data of PCV2 patients, we uncover that subclinical PCV2 reprograms the host into a striking state of non-infection, which explains its failure to respond to an initial phase of circoviral presence. A PCV2 disease signature further demonstrates that the silenced immune system associated with subclinical PCV2 becomes fully activate in PCV2 patients, triggering severe circoviral disease. Further genomic and functional analysis demonstrate STAT3 as a druggable host factor and IL-2 as a disease driver. Together, this study demonstrates the mechanistic underpinnings of clinical outcome of PCV2 infections: subclinical and clinical PCV2 display two entirely opposing transcriptomic recalibrations of lymphoid tissue.

## Introduction

Porcine circovirus type 2 (PCV2) manifests itself through a range of often devastating pathologies in swine livestock, causing severe economic losses. The most prominent disease associated with PCV2 is post-weaning multisystemic wasting syndrome (PMWS). PMWS patients exhibit progressive weight-loss, respiratory distress, pallor of skin, digestive disorders and sometimes jaundice, coinciding with pneumonia, nephritis, hepatitis and severe lymphadenopathy. Pathologic hallmarks in wasting pigs include progressive lymphocytic depletion and monocyte infiltration in lymph nodes [1], drastically compromising the immune system with often fatal outcome [2]. Although PCV2 is acknowledged as the causative agent of PMWS, PCV2 infection alone generally results in a persistent low-level replication without clinical signs [3]. In fact, PCV2 circulates endemically in pig populations as covert subclinical infections, seemingly undeterred by vaccination [4]. Pigs with PMWS however, are nearly always presented with concurrent microbial infections, suggesting a crucial role for superinfections in triggering PMWS [5]. Indeed, coinfections or other immunostimulations such as adjuvant administration were confirmed to produce PMWS in experimental models [6]. Despite two decades of intensive research, real mechanistic insights into how PCV2 achieves subclinical persistence and why certain individuals transform from subclinical PCV2 to PMWS remain unknown.

Large data sets measuring the transcriptomic architecture of biological systems combined with new mathematical and statistical models are currently revolutionizing biomedical research. Major ongoing efforts focus on identifying and genetically perturbing regulatory networks at the core of pathological processes, to enable disease outcome prediction [7,8], phenotype classification [9,10] and drug discovery [11,12]. Specifically for the field of porcine biology, many individual data sets from live animals were analyzed within the study for which they were generated, and thus, integrated analysis of the recent wealth of transcriptomic data opens opportunities for systems biologists. Here we take advantage of large volumes of porcine transcriptomic studies to create a novel gene signature collection with hundreds of gene sets characterizing *in vivo* tissue perturbation, which we subsequently interrogated against circovirus patient studies to gain mechanistic insights into the pathogenesis of host responses to PCV2 viral infection.

## Results

### PorSignDB: a gene set collection characterizing a compendium of *in vivo* transcriptomic profiles

We first created PorSignDB, a collection of gene signatures, using a systematic approach previously developed for inference of the immunologic gene signature collection ImmuneSigDB [13]. Specifically, we compiled a large gene expression compendium curated from 88 studies including 1776 unique samples. A total of 412 annotated gene sets were derived from 206 pairwise comparisons identifying genes induced and repressed in one phenotype versus another, annotated as ‘UP’ (PHENOTYPE1_VS_PHENOTYPE2_UP) and ‘DOWN’ (PHENOTYPE1_VS_PHENOTYPE2_DN) gene sets, respectively (Fig 1A). To illustrate this, an example is given for a study comparing lymph nodes of uninfected pigs versus those of pigs experimentally infected with *Salmonella enterica Typhimurium* [14]. Upregulated genes (UP gene set) are highly expressed in the uninfected phenotype, while downregulated genes (DN gene set) are highly expressed in the *Salmonella*-infected phenotype (Fig 1B). Samples were predominantly derived from real-life patients or laboratory animals (1519 *in vivo* specimens and 236 *ex vivo* samples), and additionally include some from cell cultures (21 samples). The samples were derived from a multitude of different tissues (Fig 1C) and together, they describe host responses in an entire range of biological themes, with a major part stemming from studies on microbiology, gastroenterology and the cardiovascular system (Fig 1D).

**Figure 1.**
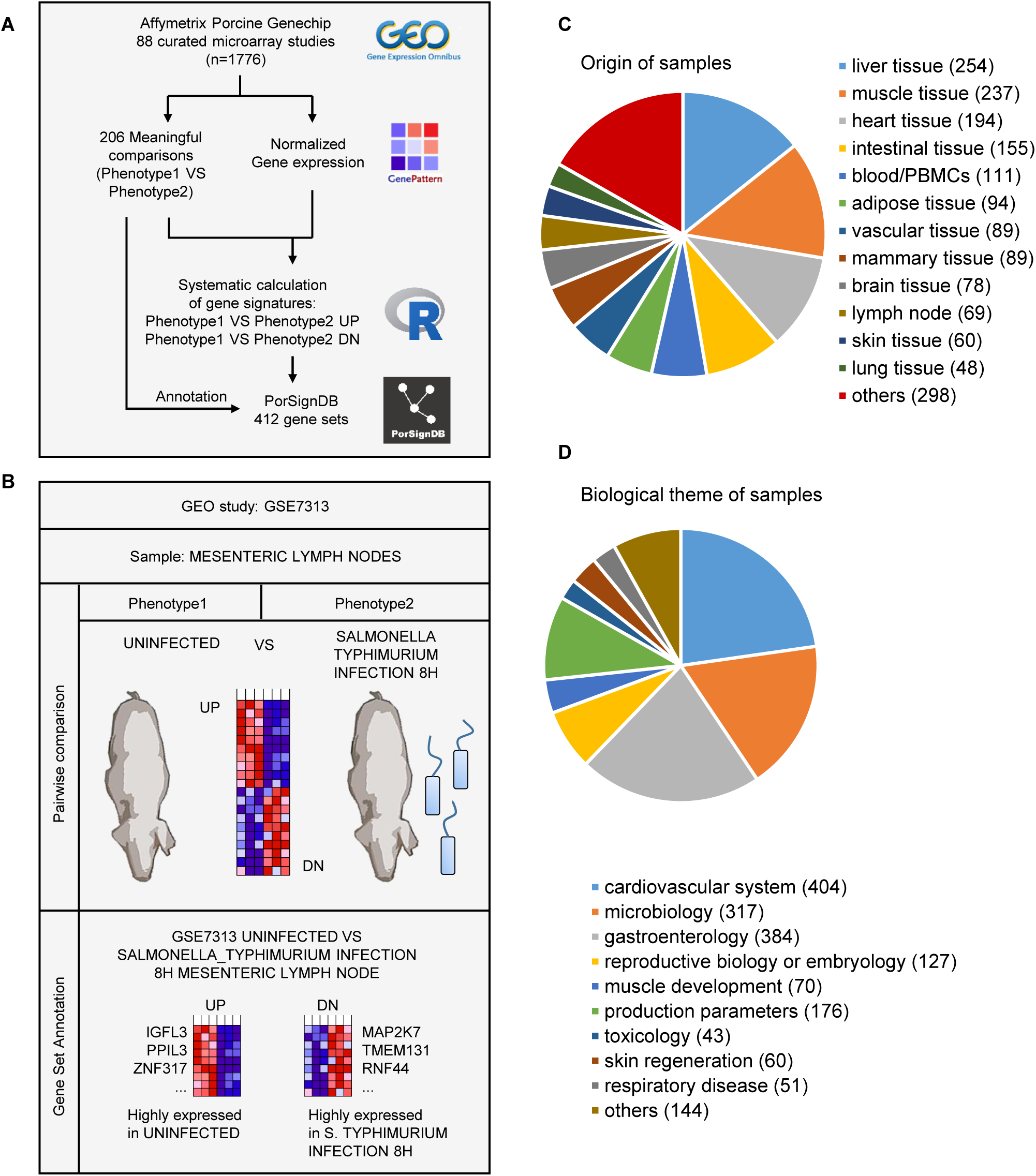
Details of PorSignDB. **A** Overview of the pipeline. 88 curated studies with data from 1776 microarrays chips were retrieved from the GEO repository. Data from each study was uniformly normalized using Genepattern, and gene expression signatures representing each phenotype of every pairwise comparison were calculated in R. Systematical annotations were added to every signature, yielding 412 gene sets. **B** Example of signature generation. GSE7313 is a study mapping transcript abundance in mesenteric lymph nodes of pigs infected with *Salmonella Typhimurium* at different time points. The first pair compares data from lymph nodes of uninfected pigs (Phenotype1) with those of pigs 8h post *S. Typhimurium* infection (Phenotype2). Significantly upregulated and downregulated genes were selected with a mutual-information based metric, respectively recapitulating highly expressed genes in the ‘uninfected’ phenotype (UP gene set), and highly expressed genes in the ‘8h post *S. Typhimurium* infection’ phenotype (DN gene set). **C** Samples were derived from a variety of different tissues, **D** covering studies in a wide range of different biological themes.

Of note, porcine genes and individual probes were mapped to *Homo sapiens* ortholog genes. Because many transcriptional programs are evolutionarily conserved, cross-species gene expression analysis can be applied successfully [15,16]. Moreover, molecular signature databases are often human-oriented, and the porcine-to-human adaptation of PorSignDB thus facilitates its application to genomic expression data of any species. The PorSignDB gene signatures are available as an online resource (http://www.vetvirology.ugent.be/PorSignDB/) and can be used by systems biologists to deconvolute cellular circuitry in health and disease. As proof of concept, we employed this gene signature collection describing host responses in a wide variety of tissues to generate new insights in the multisystemic disease associated with PCV2.

### PorSignDB reveals diametrically opposed physiological states *in vivo* in subclinical PCV2 and PMWS

We then leveraged PorSignDB to analyze a field study of pigs naturally affected with PMWS [17]. To compare transcriptomic profiles of PMWS lymph nodes with PCV2-positive but otherwise healthy lymph nodes, we tested signatures from PorSignDB for their enrichment (induced or repressed) in both classes using GSEA analysis (Fig 2A). We primarily focused on gene sets pertaining to microbiology. For robustness, we only retained signatures from pairwise comparisons in case both upregulated (PHENOTYPE1_VS_PHENOTYPE2_UP) and downregulated (PHENOTYPE1_VS_PHENOTYPE2_DN) genes are significantly enriched (FDR<0.01). For example, UP genes in splenic tissue of “control versus *Haemophilus parasuis*-infected pigs” are suppressed (Fig 2B, left heatmap first row), while DN genes are induced (Fig 2B, right heatmap first row).

**Figure 2.**
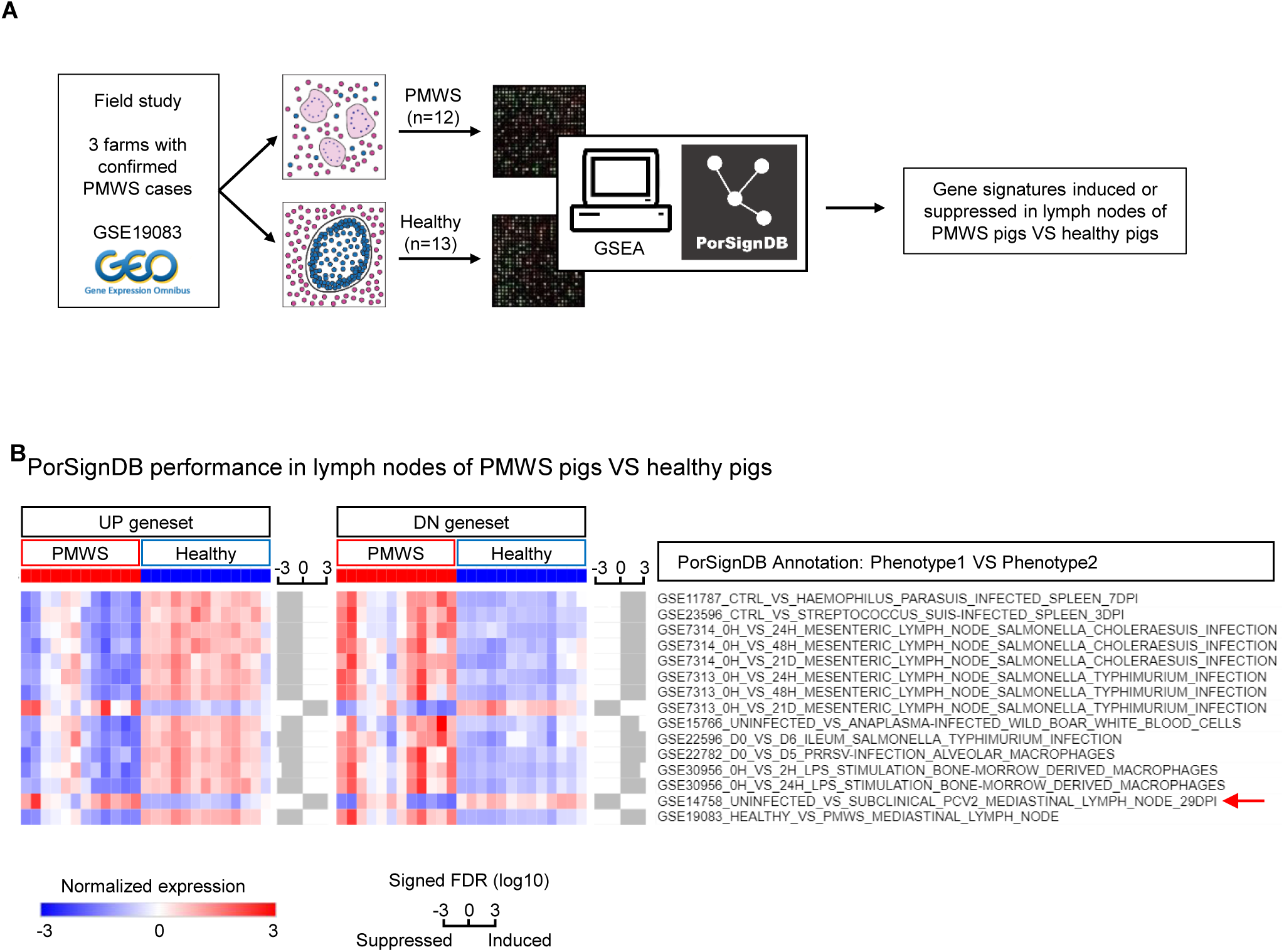
Application of PorSignDB to lymph node data originating from pig farms with naturally occurring PMWS. **A** Outline of the analysis. Data from PMWS-affected farms were retrieved from GEO. In PMWS lymph nodes, follicular structures become indistinct and B-cells and T-cells all but disappear, while infiltrating macrophages fuse into multi-nucleated giant cells. In PCV2-positive healthy lymph nodes, lymphoid structure is intact. Comparing transcriptomes of both phenotypes using GSEA displays enrichment of PorSignDB transcriptional signatures. **B** Microbiology-related PorSignDB gene set expression in lymph nodes of PMWS pigs versus Healthy pigs (FDR<0.01 and opposite expression of each pairwise phenotype). The average expression of the leading-edge genes in every gene set (genes that contribute to the enrichment) are displayed for each patient sample. Bars next to each gene set indicate the signed FDR for its enrichment in log10 scale.

Overall, this analysis reveals that upregulated genes in “control VS microbial challenge” are suppressed while downregulated genes are induced. In other words, PMWS lymph nodes display transcriptomic reprogramming consistent with tissue responses on infectious agents. This observation is supported by previous findings that naturally occurring PMWS is presented with concurrent infections [5]. Strikingly, two genomic infection signatures do not follow this pattern. First, the opposite behavior of the gene signature from *Salmonella Typhimurium* 21 days post inoculation (dpi) suggests that the *Salmonella* infection has already been cleared at this timepoint. This is indeed the case: at 21dpi the bacterial load in these mesenteric lymph nodes was reduced to undetectable levels [18]. In contrast, *S. Choleraesuis* infection was sustained at 21dpi, coinciding with persistent high bacterium abundance in mesenteric lymph nodes. Intriguingly, the second deviating gene signature originates from pigs that were subclinically infected with PCV2 (Fig 2A, arrow). Unlike *S. Typhimurium*, this cannot be explained by pathogen clearance since these experimentally PCV2-infected pigs remained viremic throughout the original study [19]. Instead, pathogen-distressed host responses appear here to be repressed in lymph nodes with low-level subclinical PCV2 replication. Hence, highly expressed genes in “uninfected VS subclinical PCV2-infected” lymph nodes are induced, while lowly expressed genes are suppressed. From this data, it can be concluded that subclinical PCV2 infection simulates pathogen-free tissue by reprogramming lymphoid tissue diametrically opposite to an ongoing infection.

Finally, the gene sets PMWS_VS_HEALTHY_UP and PMWS_VS_HEALTHY_DN serve as positive control since they were derived from the data that was queried in this instance. PorSignDB signatures from other biological themes may provide additional clues into the alterations in lymph nodes that are subject to PMWS and could be explored further (Fig S1, see also discussion).

### An immune response gene signature predicts clinical outcome of PCV2 disease

In an experimental setting, PCV2 alone does not lead to clinical signs. Additional superinfections or vaccination challenges are needed to produce PMWS [6]. Why extraneous immunostimulations trigger PMWS remains however poorly understood. A systems-level dissection of PCV2-affected lymphoid tissue may provide an explanation to this conundrum because it can determine which transcripts characterize PMWS, unbiased by previous knowledge. To this extent, the PMWS field study data was divided over a training and validation cohort, and 173 biomarker genes were selected from the training set using a leave-one-out cross validation (Fig 3A, Table S1). Together, they reveal a molecular portrait of PCV2-associated lymphoid lesions. This ‘PCV2 disease signature’ is greatly induced in the validation cohort as shown by GSEA analysis, meaning upregulation of PMWS marker genes and downregulation of Healthy marker genes (Fig 3B). Interestingly, in mediastinal lymph nodes with subclinical PCV2 at 29dpi, the disease signature is dramatically repressed when compared to lymph nodes of non-infected counterparts, showing once more that subclinical PCV2 actively suppresses the transcriptomic recalibration that goes hand in hand with PMWS. To illustrate the fidelity of the PCV2 disease signature, individual samples were classified as either PMWS or healthy with the Nearest Template Prediction algorithm [20]. All samples of the validation set were correctly assigned (FDR <0.05; Fig 3C). Furthermore, all piglets from the experimental study, either PCV2 free or with subclinical PCV2, were correctly classified as Healthy with only one sample failing to meet the <0.05 FDR threshold (Fig 3D). Furthermore, when performing a Gene Ontology overrepresentation test, the PMWS biomarker genes clearly represent the immune response, which confirms from a systems level that immune activation is a pivotal event in PMWS (Fig S2A). Of note, this gene signature performs better than an RNMI-based signature (Fig S2B-C), which is more suited for small sample sizes and was therefore applied for generating PorSignDB.

**Figure 3.**
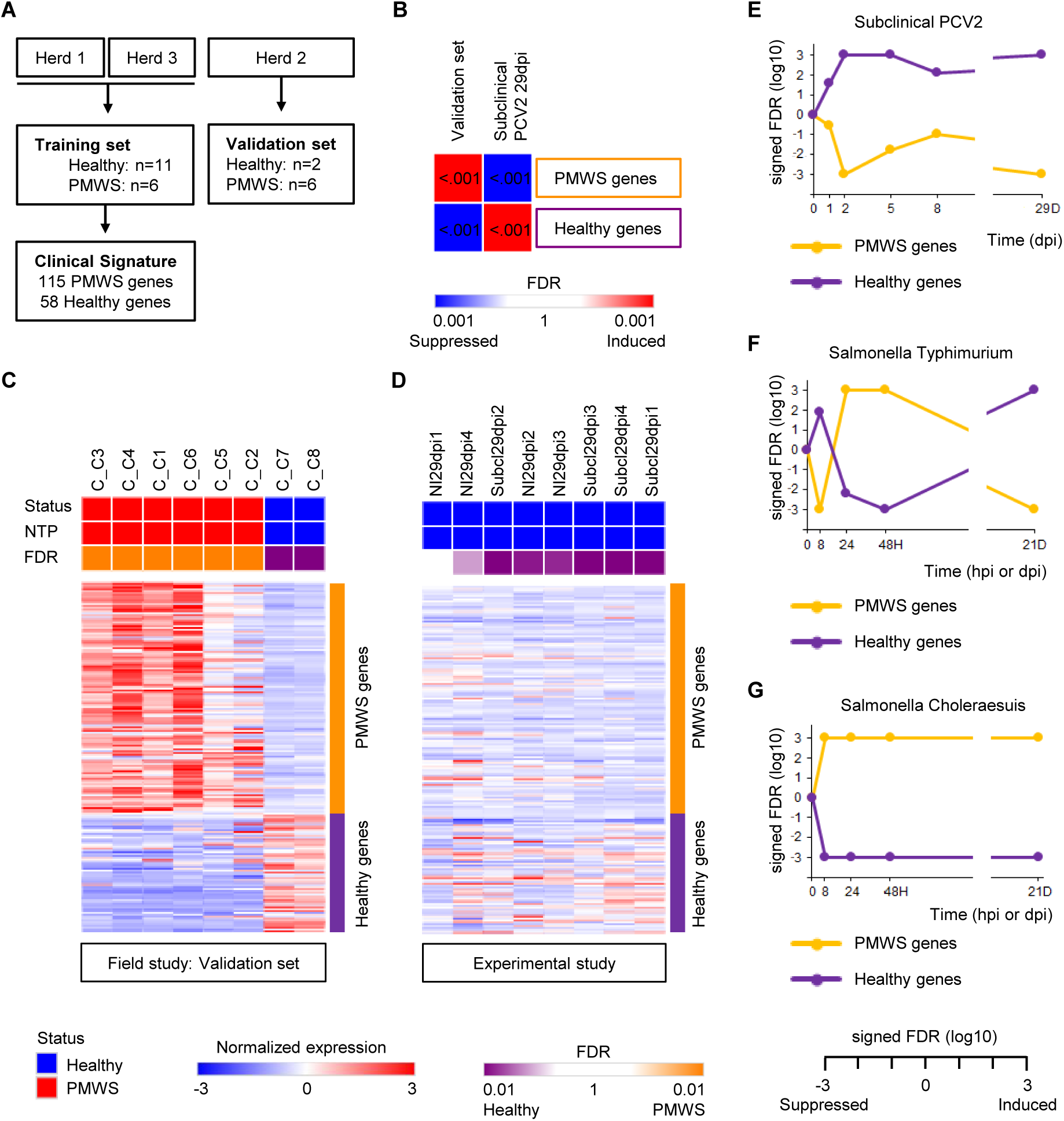
A patient-derived immune response signature predicts clinical outcome of PCV2 infection. **A** Diagram of cohort division between training and test set. A clinical PCV2 signature was calculated from the training samples and **B** tested in the validation samples by GSEA. The PCV2 disease signature was markedly induced in the validation set, and repressed in subclinical PCV2 29dpi. **C** Nearest Template Prediction of test set samples, classifying them either as healthy (blue) or PMWS (red), and **D** similarly, of the experimental subclinical infection samples at 29dpi. **E-G** Kinetics of the PCV2 disease signature upon experimental PCV2, *S. Typhimurium* and *S. Choleraesuis* infection.

Interestingly, when probing the kinetics of the PCV2 disease signature in lymph nodes of pigs experimentally infected with PCV2, *S. Typhimurium* or *S. Choleraesuis*, it is clear that these two bacterial infections promote the disease signature, while in subclinical PCV2 it is consistently suppressed (Fig 3E-G). In *S. Typhimurium* the reversal of this clinical gene signature at 21 dpi coincides with the drop of bacterial load in the mesenteric lymph nodes to almost undetectable degree. This demonstrates from a systems-approach that the infection has been virtually cleared at this time point, unlike mesenteric lymph nodes upon *S. Choleraesuis* infection, where the persistence of the signature correlates with an enduring high bacterial lymph node colonization [18].

Taken together, PCV2-induced lymphoid depletion and granulomatous inflammation in PMWS patients can be summarized in a robust gene expression signature emblematic of an enduring immune activation. Moreover, through an impartial systems-approach, we irrefutably show that a subclinical PCV2 inoculation provokes a striking suppression of the immune response.

### Functional genomics identify regulatory networks perturbations in PCV2 disease

It is becoming increasingly clear that PMWS and subclinical PCV2 represent two opposing adaptations of lymphoid tissue to circoviral infection. The former enhances immune system activation and fulminant viremia, whereas in the latter the immune system stays deafeningly silent with only a mild viremia. To understand how this tiny virus arranges this *tour de force*, the data sets covering both the PMWS field study [17] and the experimentally induced subclinical PCV2 at 29 dpi [19] were interrogated in the GSEA computational system with the innovative Hallmark gene set collection [21]. This provides a very sensitive overview of alterations in a number of key regulatory networks and signaling pathways in both PMWS patients (Fig 4, left column) and pigs with persistent subclinical PCV2 (Fig 4A, right column). In lymphoid tissue of pigs with PMWS, many of the affected transcriptional networks echo key events in PCV2-associated lymphopathology such as blatant inflammatory activity (Hallmark gene set ‘Inflammatory response’) and caspase-mediated cell death (‘Apoptosis’). Increases in gene expression mediated by p53 (’p53 pathways), reactive oxygen species (‘ROS pathway’) and NF-κB (‘TNFα signaling through NFκB’) reflect findings that PCV2 promotes p53 expression [22,23] and triggers NFκB activation through ROS [24,25] (Fig 4, left column). This analysis not only indicates critical adjustments for amplified viral replication, it also uncovers several previously unknown network modifications. These include immunological programs (‘Interferon alpha response’ and ‘Interferon gamma response’), cell signaling cascades (‘IL2-STAT5 signaling’, ‘IL6-JAK-STAT3 signaling’, ‘KRAS signaling up’) and bioenergetics (‘Glycolysis’ and ‘Hypoxia’).

**Figure 4.**
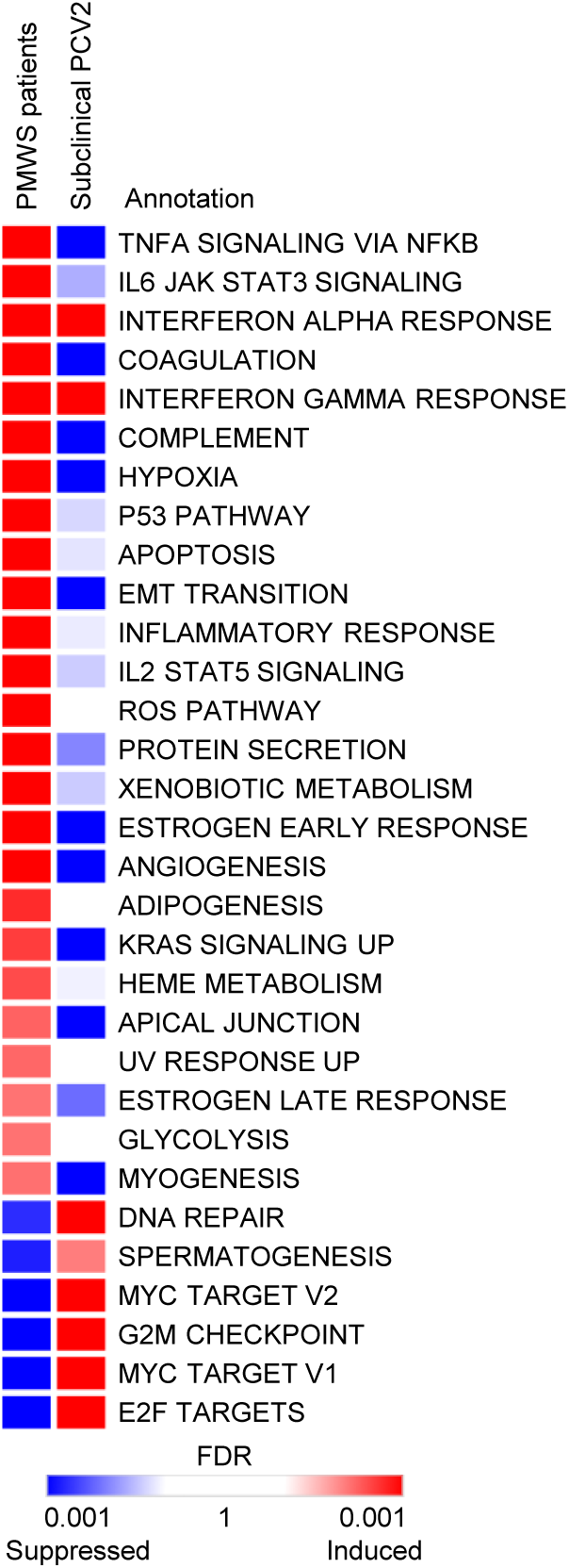
Functional genetic networks of the Hallmark gene set collection that are markedly altered in lymph nodes of pigs with PCV2. Left column: expression level in lymph nodes of PMWS patients (FDR<0.01). Right column: expression-level of these biological circuits in Subclinical PCV2 at 29dpi.

Consistent with previous results, subclinical PCV2 infection generally fails to reproduce the imbalances associated with PMWS (Fig 4, right column). Only the transcriptomic programs downstream of interferon-α and interferon-γ are in line with subclinical infections, suggesting a direct viral effect on these immunological networks. Most programs are however unaffected or opposed to the changes occurring in PMWS, reaffirming the running thread that subclinical PCV2 is unable to reprogram the circuitry to develop PMWS.

### IL-2 supplementation enables *ex vivo* modelling of PCV2 in primary porcine lymphoblasts

The transcriptional upregulation of IL-2 responsive genes in PMWS, but not in subclinical PCV2 (Fig 4A), indicates that fulminant PCV2 replication occurs in an IL-2 infused lymphoid environment. Given the pivotal role of IL-2 in activated T-cells during immune response [26], IL-2 may indeed be a crucial factor in boosting subclinical PCV2 towards PMWS. Intriguingly, the IL2-STAT5 signaling network is suppressed in subclinical PCV2, but not in *S. Choleraesuis* and *S. Typhimurium*, where there is a persistent and transient induction respectively (Fig 5A). Again, in *S. Typhimurium*, the reversal of the IL-2 signature coincides with bacterial clearance.

**Figure 5.**
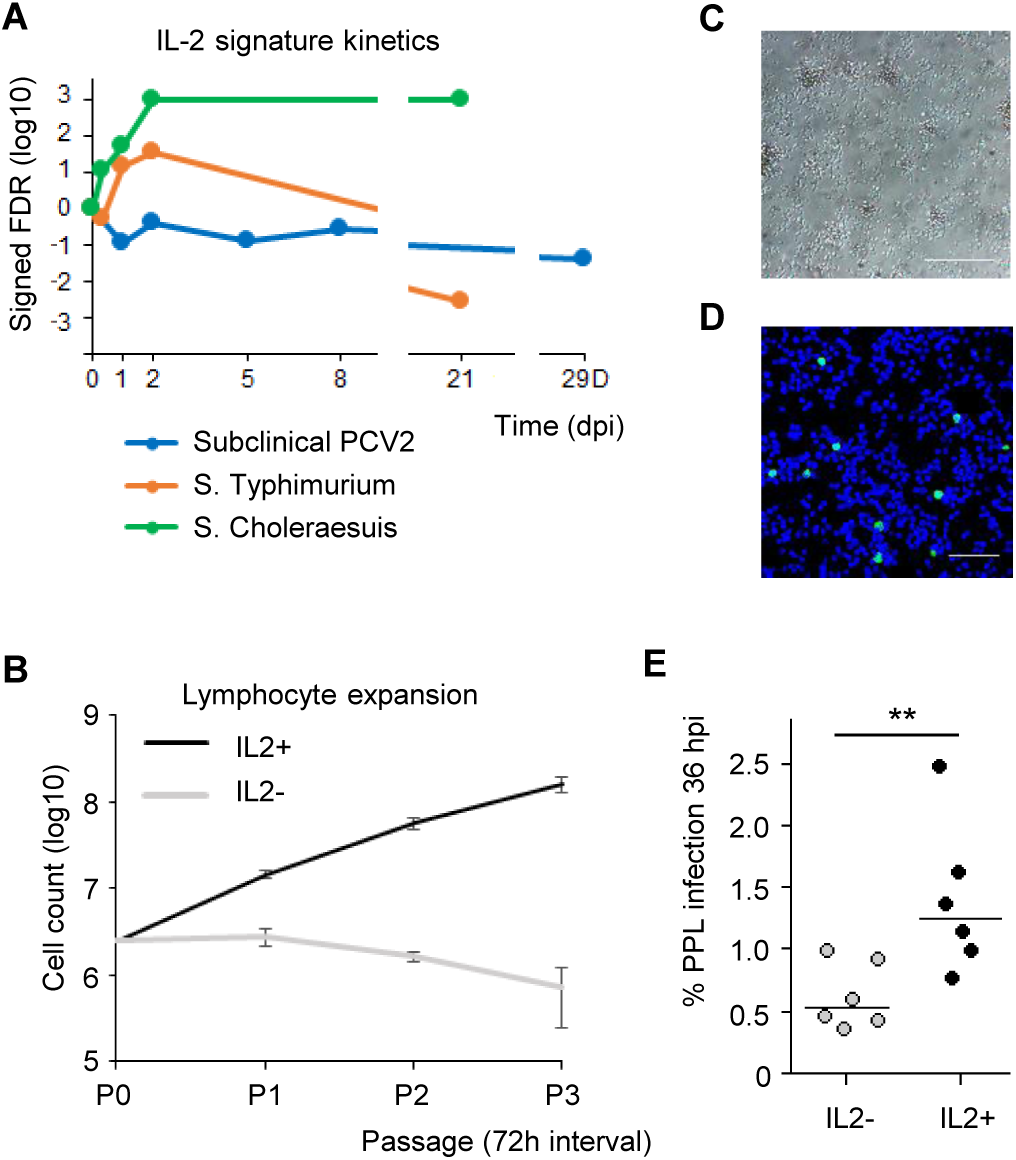
IL-2 is implicated in PCV2 disease. **A** Kinetics of IL-2 responsive gene expression (Hallmark IL2-STAT5 SIGNALING) upon three microbial infections: PCV2 (blue), *S. Typhimurium* (orange) and *S. Choleraesuis* (green). **B** IL-2 activation of freshly isolated and ConA-stimulated lymphocytes maintains exponential cellular proliferation, yielding primary porcine lymphoblast (PPL) cell strains. Means ± sd represent one experiment in triplicate (n=3). **C** Representative image of proliferating PLLs. Scale bar: 50 µm. **D** PCV2 Cap immunostaining in PLLs 36hpi. Scale bar: 100 µm. **E** IL-2 supplementation doubles PCV2 infection after a single round of replication (36 hpi). Dot blot shows six single independent experiments with horizontal line indicating median value (n=6; ^**^P<0.01, two-tailed Mann-Whitney). PPL cell strains were generated from six different individuals.

The impact of IL-2 on PCV2 replication cannot be faithfully demonstrated with traditional PK15 kidney cells. Because PCV2 has a tropism for lymphoblasts, these are the cells of choice. Our lab previously demonstrated that treatment of freshly harvested PBMCs with concanavalin A (ConA) coerces T-cells into mitosis, rendering them permissive for PCV2 [27]. Unfortunately, lymphoblast proliferation can only be maintained for a very short time after which the cells forfeit viability and die of attrition. Indeed, when isolated lymphocytes are stimulated with ConA without IL-2, these cells start suffering from apoptosis even before the first passage at 72h. However, supplementing ConA-stimulated lymphocytes with IL-2 not only mimics the PMWS microenvironment, it generates continuously expanding primary porcine lymphoblasts (PPLs; Fig 5B-C). These PPLs can be easily cultured, expanded and infected with PCV2 *ex vivo*, providing a cell culture platform amenable for studying PMWS pathogenesis (Fig 5D). To prove the beneficial effect of IL-2 on PCV2 replication, lymphocytes were freshly harvested from six individual pigs. IL-2 supplementation doubled PCV2 infection rates after 36h, a timeframe amounting to a single round of replication (Fig 5E). This demonstrates that T-cell activation by IL-2 fosters PCV2 infection and highlights the potential of PPLs for studying PMWS.

### STAT3 is a PCV2 host factor and a target for antiviral intervention

Since transcriptional networks of PMWS lymphoid tissue are subject to dramatic changes that correlate with fulminant PCV2 replication, counteracting these alterations can potentially harm the viral life cycle. When observing a fierce induction of gene expression downstream the IL6-JAK-STAT3 signaling cascade in PCV2 patients (Fig S2A), STAT3 emerges as a druggable candidate host factor. Interestingly, STAT3 is a key regulator of inflammation often exploited by viruses with pathogenic consequences [28]. In a drug assay, treatment with selective STAT3 inhibitor Cpd188 exhibits a dose-dependent effect on PCV2 infection in PPLs at 72 hpi (Fig 6A). Cell viability assay reveals no toxicity, excluding non-specific adverse effects of the compound on infection (Fig 6B). Chemical inhibition also displays a dose-dependent effect on PCV2 infection in PK15 cells (Fig S2B-D). Thus, robust expression of STAT3 responsive genes are critical for PCV2, and hampering STAT3 activity represents an antiviral strategy (Fig 6C).

**Figure 6.**
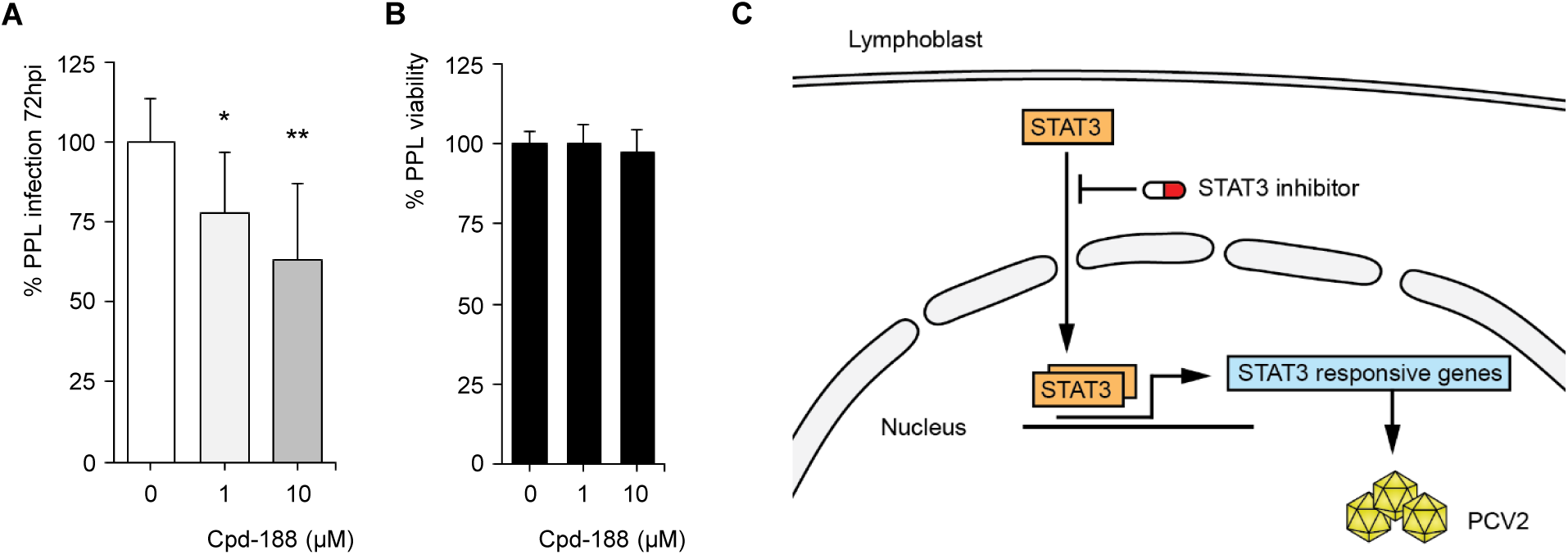
STAT3 is a PCV2 host factor. **A** STAT3-specific inhibitor Cpd188 impairs infection in PPLs. Means ± sd represent three independent experiments in triplicate (n=9; ^*^P<0.05, ^**^P<0.01, two-tailed Mann-Whitney). **B** MTT lymphoblast viability assay of Cpd188 treatment. Means ± sd are shown for three experiments in quintuplicate (n=15). **C** Cartoon outlining STAT3 as a drugable host factor for PCV2 in lymphoblasts.

### A paracrine macrophage-lymphoblast communication axis exacerbates PCV2 infection

Finally, the PMWS field study dataset was queried in GSEA with ImmuneSigDB’s immunological gene signatures [13]. This revealed a striking suppression of lymphocyte gene expression and powerful induction of signatures from monocytes and other myeloid cells (Fig 7A, Table S2), reflecting the loss of lymphocytes and histiocytic replacement in PMWS lymph nodes. This raises the question to what extent infiltrating monocytes affect PCV2 replication. After maturation into macrophages, they may either dampen infection by destroying viral particles, or promote PCV2 in a paracrine fashion by releasing pro-inflammatory cytokines. To test the effect of intercellular communication between macrophages and lymphocytes, a co-culture experiment was set up. PCV2-infected PPLs were seeded in a porous insert, physically separated from a lower compartment with primary porcine macrophages (Fig 7B). The latter were challenged with Porcine Reproductive and Respiratory Syndrome Virus (PRRSV), a virus that can experimentally trigger PMWS [6] (Fig 7C). The presence of non-infected macrophages had no significant effect on PCV2 lymphoblast infection levels, but when co-cultured with PRRSV-infected macrophages, a significant and consistent increase in PCV2 infection could be discerned (Fig 7D). This demonstrates the existence of a previously unknown axis of intercellular communication between macrophages and lymphoblasts exacerbating PCV2 replication.

**Figure 7.**
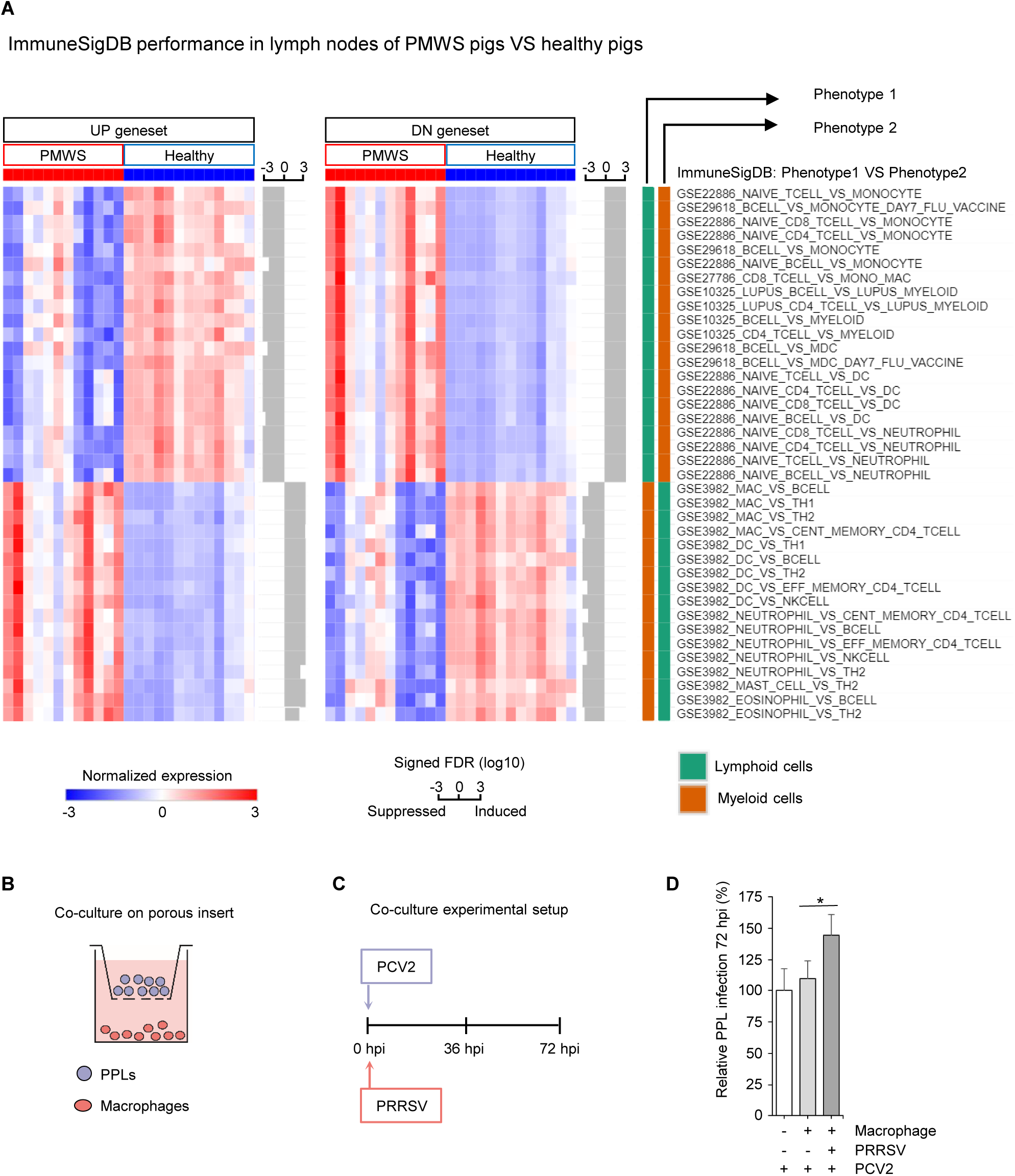
Superinfection increases PCV2 replication through a macrophage-lymphoblast paracrine signaling axis. **A** ImmuneSigDB gene set expression in the PMWS field study (FDR<0.01 and opposite expression of each pairwise phenotype). The average expression of the leading-edge genes in every gene set (genes that contribute to the enrichment) are displayed for each patient sample. Bars next to each gene set indicate the signed FDR for its enrichment in log10 scale. PMWS versus healthy lymph node comparison displays a dramatic repression of lymphocyte gene expression signatures, and induction of myeloid cell signatures. **B** Experimental set-up of PPL-macrophage co-culture system mimicking PMWS lymph nodes. **C** PCV2-inoculated PPLs were seeded on a porous insert with macrophages at the bottom of the well. Macrophages were additionally challenged with PRRSV at 0h. **D** Relative PPL infection levels at 72hpi. Means ± sd represent two independent experiments in triplicate (n=6; ^*^P<0.05, two-tailed Mann-Whitney).

## Discussion

Systems biology approaches aim to model regulatory networks from genome-wide transcriptional profiles as a large interconnected switchboard where many small inputs combine to execute large programs. In response to extra- or intracellular stimuli, cells will modify their circuitry in an orchestrated effort to adapt to their environment. These days, mapping genome-wide transcripts for biological network analysis has become fast, cheap and easy. Especially when deposited in public databases, it provides an ever-growing library of transcriptomes. Here we unlock the potential of porcine microarray studies by turning it into an atlas of tissue transcripts for bioinformatical analysis, extending the MSigDB database with *in vivo* derived profiles [29]. Prior mapping of porcine microarray probes to human orthologs facilitates its application to any mammalian gene expression data set.

PorSignDB is especially convenient for delineating which physiological state one’s samples of interest resemble, generating useful hypotheses in the process. When applied to PCV2 patient data, it elegantly exposes the transcriptomic bedrock underlying circoviral persistence: PCV2 reprograms lymph node circuitry into a state of non-infection as a strategy for establishing covert chronic infection. We hereby postulate the mechanics of how PCV2 operates in swine livestock. In an initial phase, PCV2 functionally silences the immune response, delaying or even completely abrogating an adaptive antibody response [30]. This recalibration allows only low-level PCV2 replication but does result in viral persistence without clinical signs. Only when PCV2’s input on host transcriptomics is overturned by a severe superinfection are inflammatory networks and STAT3 responsive gene expression engaged. This immune activation infuses lymphoid tissue with IL-2 to activate T-cells into lymphoblasts, triggering fulminant circoviral replication and widespread lymphocyte apoptosis. When macrophages rush in to help, their paracrine signaling exacerbates PCV2 replication even further. Moreover, PCV2 capsids resist destruction by macrophages, which fuse together into histiocytic giant cells in an ineffective last-ditch effort to stem the infection. PMWS is thus the end-stage of a lethal viral lymph node disease, where germinal centers have collapsed and functional parenchyma is replaced by macrophages, leading to a structural immune deficiency.

Confronted with the evidence that subclinical PCV2 and PMWS are two different host reactions to PCV2, we deem it important to discriminate between these two phenotypes of ‘PCV2 infection’. Treating them as a single entity will only result in conflicting data. As an example, this integrative transcriptional analysis resolves the long-standing dichotomy in PMWS pathology of whether or not apoptosis is implicated in lymphoid depletion *in vivo* [31–33]. In lymphoid tissue with low-level replication, it is not. On the other hand, in lymph nodes collapsing under PCV2, genes mediating apoptosis are in full force (Fig 4).

Another example of PorSignDB generating intriguing hypotheses, is that weaned gut gene expression signatures are induced in clinical PCV2, while intestinal signatures of suckling piglets are suppressed (Fig S1). It suggests that as long as intestinal tissue is protected by maternal antibodies, progression to PMWS is obstructed. On the other hand, when weaned, naive intestinal tissue makes immunological contact with pathogens, producing a microenvironment that reflects PMWS and hence, can promote PCV2.

Finally, the pronounced IL-2 signature in clinical PCV2 inspired the establishment of primary lymphoblast strains. They can be easily expanded and stored in liquid nitrogen, and display excellent post-thaw survival. Unlike PK-15 cells, they can be harvested from different individuals or breeds, providing a new and valuable tool for studying the long-suspected impact of genetic background on PCV2 disease [34,35].

In conclusion, we here solve a long-standing enigma of how PCV2 establishes subclinical persistence, and how it switches to clinical disease. Upon infection, host tissue is instructed to act as if the pathogen is absent, allowing PCV2 to replicate covertly at modest rates. Whenever an individual falls victim to a stimulus that rewires the transcriptional circuitry into immune activation, PCV2 replicates frantically and overwhelms the host. Given its limited coding capacity, PCV2 cannot manage it alone but depends on superinfections to recalibrate the host. This elegantly explains how PCV2 circulates in pig farms, and settles the controversies that have haunted PCV2 pathologists.

## Materials and methods

### Generating PorSignDB

Raw Affymetrix Porcine Genechip data were retrieved from NCBI GEO (http://www.ncbi.nlm.nih.gov/geo/). Data covering pooled samples or lacking publication on Pubmed were discarded, as were studies with <2 samples per phenotype. Quantile normalized expression data was generated from CEL files using the ExpressionFileCreator module on Genepattern [36]. Affymetrix probe set identifiers were mapped to Homo sapiens gene symbols as previously described [37] with Refseq and Uniprot identifiers were changed into corresponding gene symbols. Early transcriptional responses (<30 mins) and comparisons between breeds or tissue types were ignored. If controls were unavailable for temporal studies, comparisons were made with t=0. For signature generation, the ImmuneSigDB recipe [13] was followed. Briefly, genes were correlated to a target profile and ranked using the RNMI metric [38]. Top and bottom ranked genes with an FDR <0.01 in a permutation test were included in two gene sets, with maximally 200 genes each, yielding PHENOTYPE1_VS_PHENOTYPE2_UP" and “PHENOTYPE1_VS_PHENOTYPE2_DN”.

### PCV2 disease signature and phenotype classification

Biomarker genes were calculated from data of a field study covering three different cohorts [17], according to a previously described method [7] with minor modifications. Cohorts were divided over a training set (n=17) and a validation set (n=8). Marker genes were ranked in the training set using signal-to-noise ratio (S2NR), with standard deviations adjusted to minimally 0.2*mean. In a subsequent leave-one-out cross validation, a single sample was left out and a permutation test was performed on the remaining samples. Only genes with p<0.05 in every iterative leave-one-out trial were included in the signature. For phenotype classification, the NTP algorithm [20] was employed with S2NR as weights.

#### Cells, virus and reagents

PK15 kidney cells were a kind gift of Gordon Allan, Queen’s University, Belfast, UK. PK15 culture conditions were described earlier [39]. To generate PPLs, PBMCs were isolated as described previously (Lefebvre *et al.*, 2008b). After adhering of monocytes to a plastic culture flask, lymphocytes in suspension were pelleted, resuspended in culture medium supplemented with 5 µg/ml ConA (Sigma) and 50 µM β-mercaptoethanol (Gibco). After three days, cells were pelleted, washed with RPMI (Gibco), and resuspended in culture medium supplemented with 100 U/ml human recombinant IL-2 (NIH) and 50 µM β-mercaptoethanol. Porcine alveolar macrophages were isolated as described [40]. Animal procedures were approved by Ghent University ethical committee EC2013/97. PCV2 strains 1121 and Stoon1010 were described previously [41]. PRRSV Lelystad strain was described earlier [40]. Cpd188 was ordered from Merck Millipore.

#### Experimental infection and immunostaining

PK-15 and PPLs were inoculated with PCV2 1121 at 0.1 TCID_50_/cell for 1h, washed and further incubated in culture medium for 36h. For Cpd188 experiments, cells were pre-incubated for 1 hour with Cpd188 (Merck Millipore) dissolved in 0.25 % DMSO. Subsequently, cells were inoculated with PCV2 at 0.1 TCID_50_/cell for 1h, washed and incubated for 72h. For co-culture, PPLs and macrophages were inoculated at 0.5 TCID_50_/cell for 1h with PCV2 Stoon1010 and PRRSV respectively, washed and incubated for 72h. PCV2 capsid immunostaining with mAb 38C1 was described earlier (Huang *et al.*, 2015).

## Acknowledgements

We thank Joaquím Segalés and Lana T. Fernandes for sharing clinical data, Hussein El-Saghire, Gerben Menschaert and Eloi E.R. Verrier for helpful discussions, and Carine Boone for technical assistance.

## Supporting Information Legends

**Figure S1.**
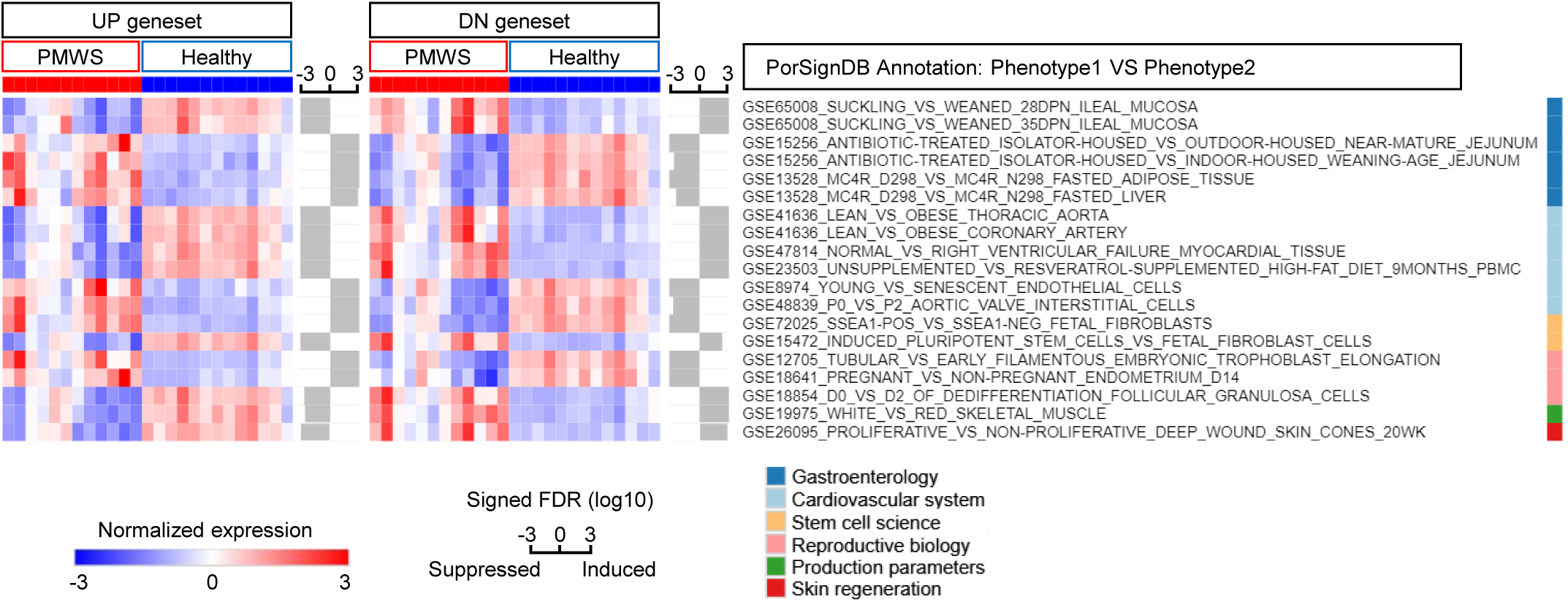
PorSignDB performance in lymph nodes of PMWS pigs VS healthy pigs. Figure displays enriched PorSignDB gene sets in the PMWS study pertaining to biological themes other than microbiology.

**Figure S2.**
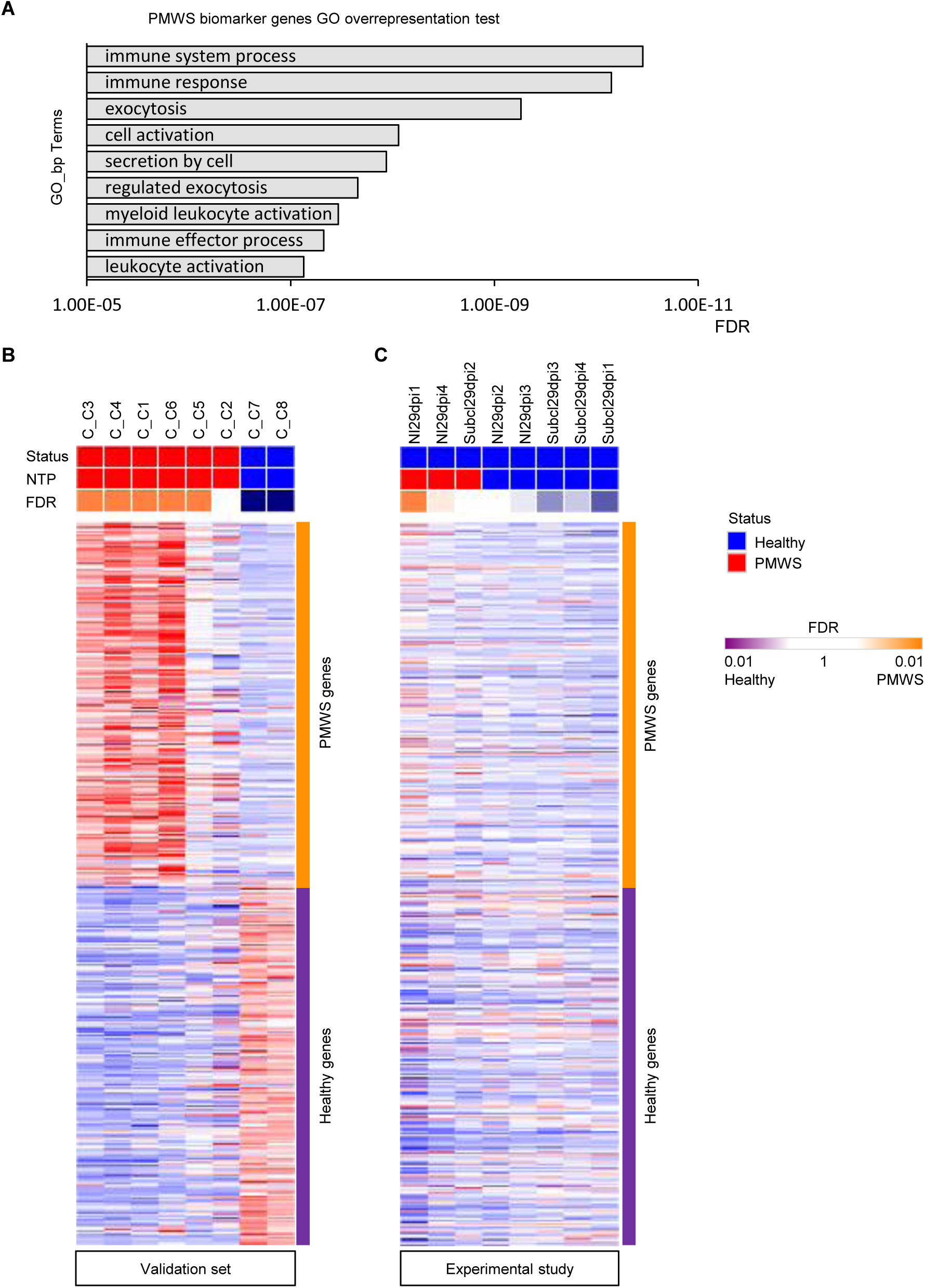
PMWS biomarker genes annotation and performance of an alternative clinical disease signature. **A** Gene ontology (GO) terms overrepresentation test of PMWS biomarker genes. **B** Nearest Template Prediction of test set samples using an alternative clincal gene signature based on the RNMI metric **C** and similarly, of the experimental subclinical infection samples at 29dpi.

**Figure S3.**
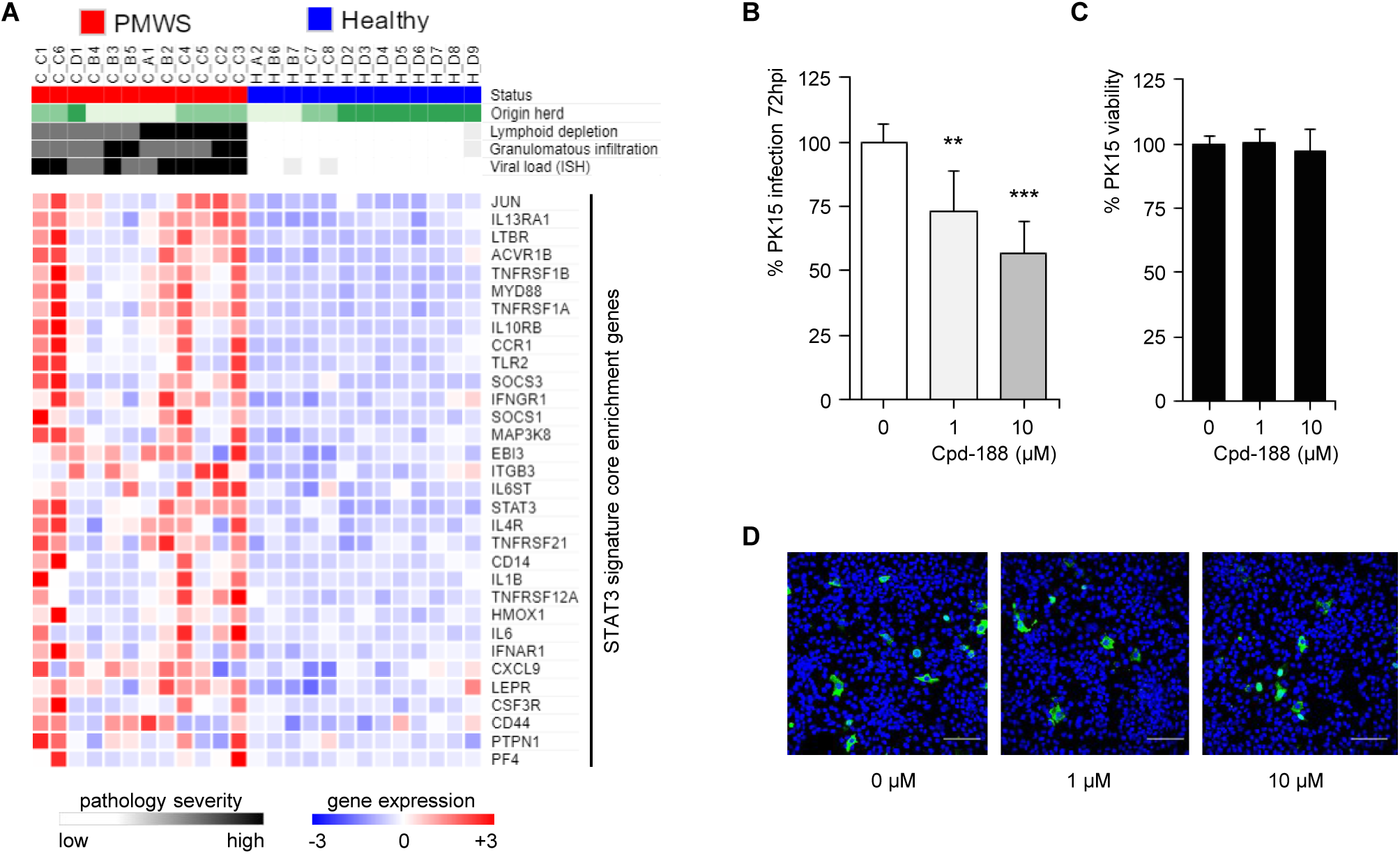
STAT3 is a host factor in PCV2 disease. **A** Core genes responsible for the STAT3 signature enrichment score. **B** STAT3-specific inhibitor cpd-188 impairs infection in PK-15 cells. Means ± sd represents three independent experiments in triplicate (n=9; ^**^P<0.01, ^***^P<0.001, Mann-Whitney U-Test). **C** MTT cell viability assay of cpd-188 treatment in PK-15 cells. Means ± sd are shown for three independent experiments in quintuplicate (n=15). **D** Infection assessment by PCV2 capsid immunostaining, representative figures for each treatment. Scale bar: 100 µm.

**Table S1** Complete list of PCV2 disease signature biomarker genes.

**Table S2** ImmuneSigDB analysis of the PMWS field study dataset.

